# Identification of Neural Crest and Neural Crest-Derived Cancer Cell Invasion and Migration Genes Using High-throughput Screening and Deep Attention Networks

**DOI:** 10.1101/2024.03.07.583913

**Authors:** JC Kasemeier-Kulesa, S Martina Perez, RE Baker, PM Kulesa

## Abstract

**Background:** Cell migration and invasion are well-coordinated processes in development and disease but remain poorly understood. We previously showed that highly migratory neural crest (NC) cells share a 45-gene panel with other cell invasion phenomena, including cancer. To identify critical genes of the 45-gene panel, we performed a high-throughput siRNA screen and used statistical and deep learning methods to compare NC- versus non-NC-derived human cell lines.

**Results:** We find 14 out of 45 genes significantly reduces c8161 melanoma cell migration; only 4 are shared with HT1080 fibrosarcoma cells (BMP4, ITGB1, KCNE3, RASGRP1). Deep learning attention network analysis identified distinct cell-cell interaction patterns and significant alterations after BMP4 or RASGRP1 knockdown in c8161 cells. Addition of recombinant proteins to the culture media identified 5 out of the 10 known secreted molecules stimulate c8161 cell migration, including BMP4. BMP4 siRNA knockdown inhibited c8161 cell invasion *in vivo* and *in vitro*; however, its addition to the culture media rescued c8161 cell invasion.

**Conclusion:** A high-throughput screen and deep learning rapidly distilled a 45-gene panel to a small subset of genes that appear critical to melanoma cell invasion and warrant deeper *in vivo* functional analysis for their role in driving the neural crest.

## INTRODUCTION

Cell migration and invasion are well-coordinated processes in early development, cancer, the immune response, and wound healing that, if uncontrolled, can lead to severe birth defects and life-threatening disease. The neural crest (NC) serves as a model for cell invasion and collective migration during vertebrate development (Szabo and Mayor, 2016, 2019; Giniunaite et al., 2019) and for several invasive human mesenchymal cancers, including NC-derived melanoma (Friedl et al., 2004; Friedl and Alexander, 2011; Kulesa et al., 2006). NC cells are essential to vertebrate embryogenesis, traveling long distances after exiting the dorsal neural tube to contribute to nearly every peripheral organ (LeDouarin and Kalcheim, 1999). Although NC cells emerge all along the anteroposterior axis and are sculpted into discrete streams that invade immature extracellular matrix (ECM) and loose mesoderm, the molecular mechanisms that underlie invasion and collective cell migration remain unclear. Mistakes in NC cell migration led to birth defects, termed neurocristopathies, that include shortened lower jaw and cleft lip and palate in the head, and failure to reach the end of the embryonic gut or Hirschsprung’s disease (Vega-Lopez et al., 2018; Fitriasari and Trainor, 2023). Therefore, by leveraging the accessibility of the neural crest to advances in spatial gene profiling and their ancestral relationship to melanoma, identification and functional analysis of a critical set of cell invasion genes have the potential to inform strategies to repair human neurocristopathies and control cancer metastasis.

Single cell transcriptome analyses have made clear that there is tremendous spatial molecular heterogeneity within a discrete NC cell migratory stream (Morrison et al., 2017; 2021; Tatarakis et al., 2021). In the chick head, isolation and profiling of cranial NC cells at distinct, progressive time points during migration within the second branchial arch (BA2) stream revealed that leader NC cells have a novel transcriptional signature that is consistent and enriched for approximately 900 genes (Morrison et al., 2017). Subsequent study of the cellular landscape of the first four chick cranial to cardiac branchial arches (BA1-4) using label-free, unsorted single-cell RNA sequencing identified 266 invasion genes out of the original list (>900 genes; Morrison et al., 2017) common to all four BA1-4 NC cell migratory streams (Morrison et al., 2021). In support of transcriptional differences within NC cell migratory streams throughout the embryo, single cell RNA-seq analysis of isolated leader and follower enteric neural crest-derived cells (ENCDCs) showed that leader ENCDCs are transcriptionally distinct from followers, exhibiting altered expression of ECM and cytoskeletal genes consistent with an invasive phenotype (Stavely et al., 2023). Wavefront-ENCDCs also lacked expression of genes related to neuronal or glial maturation, present in follower cells (Stavely et al., 2023) and support recent results showing zebrafish NC cell lineage decisions are made during mid-migration rather than at the end target (Tatarakis et al., 2021). Although these transcriptional data offer an opportunity to better understand NC cell invasion and collective cell migration mechanisms, the future studies typically to determine the function of individual genes derived from large-scale transcriptional analyses are time-prohibitive and impractical.

To address this challenge, high-throughput screening offers a rapid means to analyze large transcriptional data with an initial prioritization relevant to cell migration and invasion. Unfortunately, dissection and isolation of large numbers of embryonic NC cells in any current model organism with the purpose to seed multiple, multi-well high-throughput assays are extremely labor intensive and unreasonable. To overcome this, the readily available NC-derived human metastatic melanoma c8161 cell line which displays similar behaviors to embryonic NC cells (Kulesa et al., 2006; Bailey et al., 2012; Bailey and Kulesa, 2014) including aggressive invasion and collective cell migration can instead be used. Knockdown of individual genes using siRNA transfection (Table 1) in cell culture offers a straightforward means to molecularly perturb gene function and, when combined with a high-throughput assay and time-lapse confocal microscopy, permits quantification and comparison of changes in cell migratory behaviors and invasion.

**Table 1:** siRNAs used on 45-gene panel list.

| C8161+HT1080<br>Inhibitors | C8161<br>Specific<br>Inhibitor | HT1080<br>Specific<br>Inhibitors | C8161<br>Agonists |
| --- | --- | --- | --- |
| ARPC3 | BMP4 | CA2 | KITLG |
| PRKCQ | IFI30 | DPYD | RASGRP3 |
| RASGRP3 | IL13RA1 | HSPB1 | VEGFC |
| CREB3L2 | INHBA | INHBA |  |
| ITGB1 | JUN | MMP11 |  |
| UNC5B | KYNU |  |  |
|  | NEXN |  |  |
|  | NRP1 |  |  |
|  | RAC2 |  |  |
|  | RASGRP1 |  |  |
|  | SERPINI1 |  |  |
|  | SH3TC1 |  |  |
|  | TUBB6 |  |  |
|  | UPP1 |  |  |
|  | VIM |  |  |

Collective cell migration and invasion are critical in a wide range of diverse biological phenomena, including embryogenesis, wound repair, the immune response, and cancer metastasis. Since the neural crest are widely utilized as a model system to study collective cell migration and invasion mechanisms, a list of 34 published gene signatures was curated from other invasive cell types and compared to the embryonic NC cell invasion signature (Morrison et al., 2021). This comparison revealed a subset of 45 out of greater than 900 genes of the chick leader cranial NC cell invasion signature in common with two or more cell invasion signatures from this list (Morrison et al., 2021). This condensed signature of 45 genes, although not encapsulating all genes necessary for cell migratory and invasive characteristics, is manageable for a high-throughput siRNA screening assay to prioritize a smaller subset of genes for single gene *in vivo* functional studies of NC cell migration. Results of such a screening assay will also help to determine whether these 45 genes have a conserved function in non-NC-derived cell types and more broadly to other cell migration and invasion phenomena.

In this study, we compared changes in cell dynamics in human metastatic melanoma c8161 versus human fibrosarcoma HT1080 cell lines after siRNA gene knockdown. Using a plugged plate assay in which cells are seeded around a rubber stopper, open space closure rates, as well as the dynamics of edge (leader) and bulk (follower) cells, were measured after siRNA gene knockdown and rubber stopper removal as well as the dynamics of edge (leader) and bulk (follower) cells into open free space to determine significant changes in cell invasion and collective cell migration. We then used deep learning attention network analysis of large-scale cell trajectory data generated from each of the gene knockdowns to identify alterations in cell neighbor interactions. Separately, we performed an analysis of the known secreted molecules of the 45-gene panel by comparing changes in c8161 cell migration after addition of the recombinant proteins in the culture media. Together, these experiments identify a subset of the 45 gene invasion signature that affects melanoma cell migration and warrants further in-depth analysis.

## RESULTS

### A rubber stopper high-throughput screening assay avoided disruption of cell membranes and provided a uniform intact circular migratory wavefront for analysis of open free space invasion

To systematically test our novel 45 gene invasion signature derived from a comparison between the embryonic cranial NC cells and a broad range of other cell invasion phenomena (Fig. 1A), we used a high- throughput rubber stopper-based screening assay to study human c8161 melanoma versus HT1080 fibrosarcoma cell lines (Fig. 1B). Each cell line was transfected with siRNAs to knockdown single genes in duplicate from the 45-gene panel prior to seeding cells in rubber stopper-plugged plates (along with water and negative siRNAs for controls). After seeding and adherence of cells in each well, the plugs were removed from each well and cell trajectories were collected from 24hr time-lapse confocal imaging sessions (Fig. 1B, C). Open free space was defined as the area created by removal of the rubber stopper plug in each well. In addition, removal of the rubber stopper did not disrupt cell integrity and provided a uniform circular wavefront (Fig. 1C), in contrast to the typical wound closure associated with scratch assays. Cell dynamics data was analyzed using traditional metrics (for example, mean-square displacement and directionality) as well as using deep learning networks to study changes in cell-neighbor relationships after siRNA knockdown of each of our 45-gene panel (Fig. 1D). Lastly, we analyzed the 45-gene panel for upstream transcription factors (Fig. 1E).

**Figure 1:**
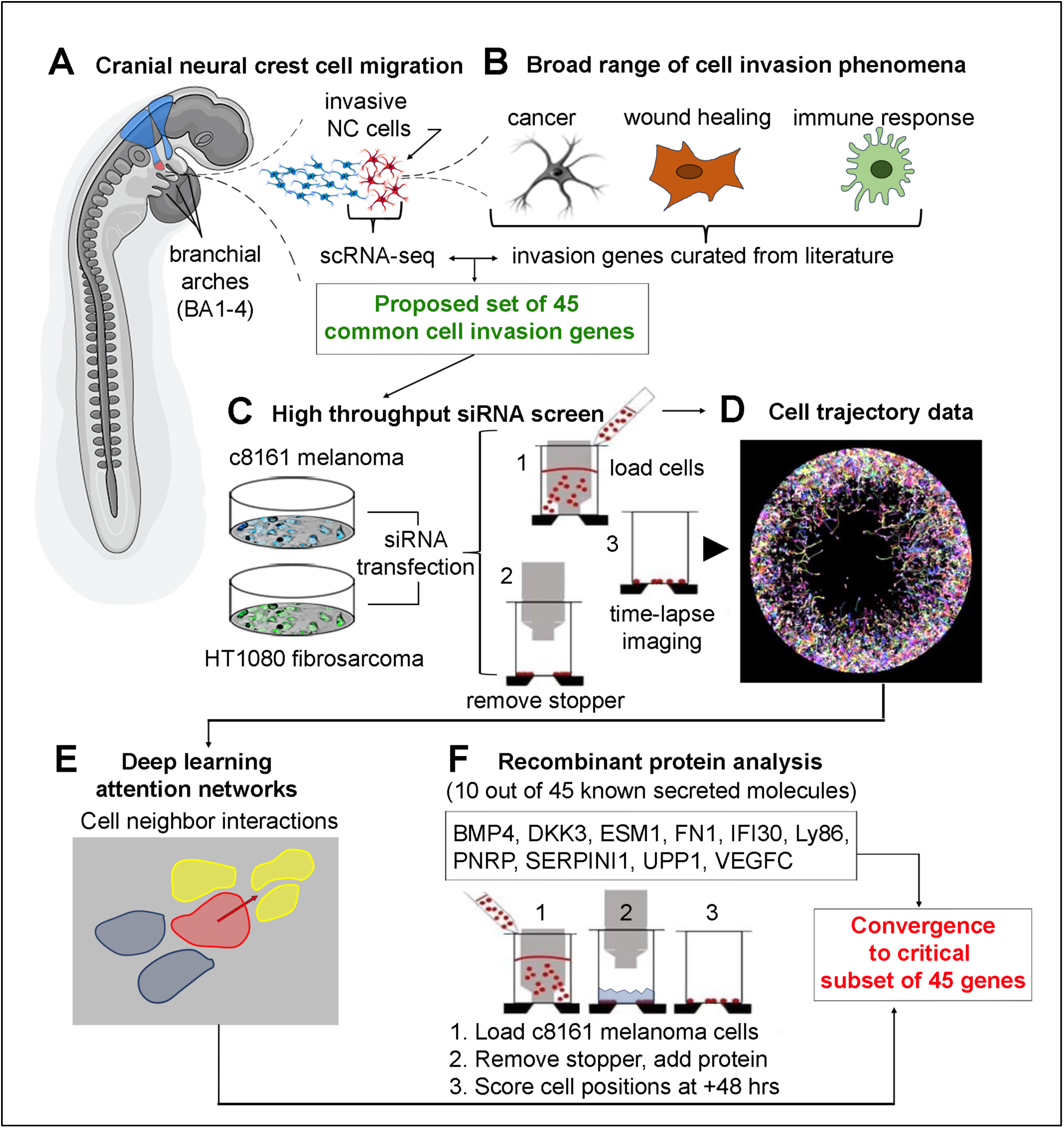
Schematic of integrated experimental and computational workflow of high throughput screen assay and cell behavior analytics. (A) Cranial neural crest (NC) cell migration in the vertebrate head in discrete streams with a migratory wavefront (red) and follower NC cells (blue). (A-B) Comparison of genes enhanced within NC cells at the migratory wavefront and published invasion signatures from a wide range of other cell invasion phenomena. (C) High-throughput screen compares changes in migration and invasion behaviors of c8161 melanoma versus HT1080 cells after gene knockdown, using a plugged plate assay and confocal time-lapse imaging to (D) generate dynamic cell trajectory data. Scoring of cell positions at +48 hours after plug removal (static analysis not shown) and (E) deep learning network analysis to analyze changes in cell-cell neighbor interactions. (F) Recombinant protein analysis performed separately and static analysis to determine changes in open space closure in the presence of 10 out of 45-gene panel known secreted molecules, using same plugged plate assay.

### The invasion of c8161 melanoma cells into open free space is significantly affected by knockdown of 14 out of 45 genes that primarily affect cell motility rather than proliferation

As an initial screening examination to assess which siRNA knockdowns were associated with dynamic changes in human cancer cell behaviors during closure of the free space, we performed a statistical analysis of free space area and cell count fold changes (Fig. 2). For each well, we computed z-statistics for the free space area and cell count fold change. We present gene knockdowns that significantly affected free space closure defects and find 14 out of 45 genes included in the screen resulted in significant free space closure in the c8161 melanoma cells (Fig. 2A). BMP4, ITGB1, IFI30, KCNE3, PRKCQ, and RASGRP1 ranked in the top six out of 14 (Fig. 2A). Strikingly, in the HT1080 fibrosarcoma cells only four out of 14 genes exhibited significantly reduced free space closure (BMP4, ITGB1, KCNE3, and RASGRP1; highlighted in orange and shared with the top six genes of the c8161 melanoma cells in Fig. 2A).

**Figure 2:**
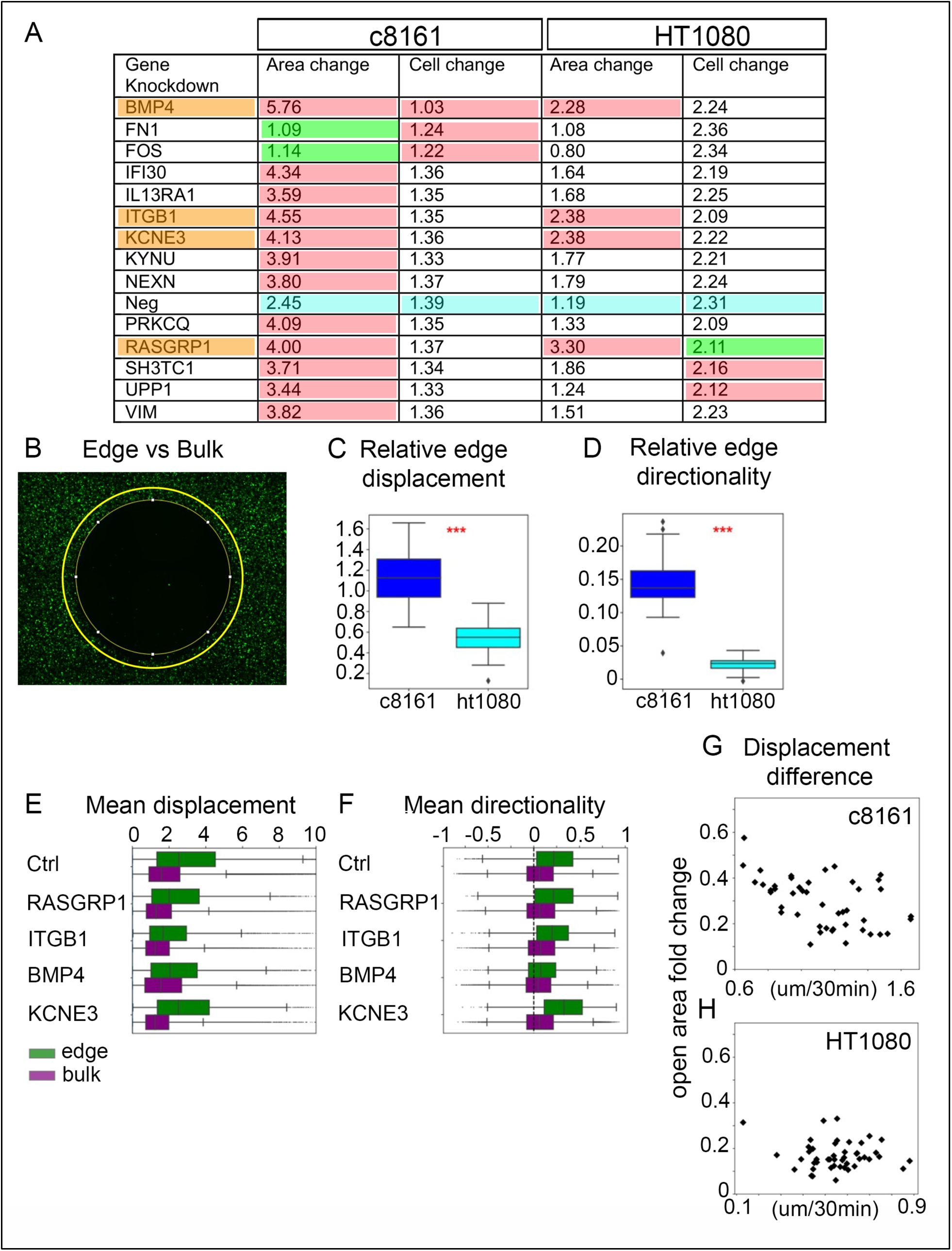
Quantitative metrics of open area closure and migration. (A) Table of genes resulting in significantly reduced wound closure in the c8161 cell line with fold changes for wound area and cell count fold changes. (B) TrackMate tracks overlayed for cells in the edge region. (C) relative edge displacement across all experiments using c8161 and ht1080 cells. (D) Relative edge directionalities across all experiments using c8161 and HT1080 cells. (E) Mean displacement in edge and bulk regions in the c8161 cell line. (F) Mean directionality in edge and bulk regions in the c8161 cell line. (G) Displacement difference vs wound closure for all considered c8161 biological replicates. (H) Directionality difference vs wound closure for all considered HT1080 biological replicates.

To better understand the interplay between cell proliferation and cell motility during free space closure, we compared the two characteristics. In the c8161 melanoma cells, only BMP4 knockdown inhibited free space closure in conjunction with significantly reduced cell proliferation (Fig. 2A; first row, shaded in red). Other knockdowns of the 14 out of 45 genes that exhibited a reduced rate of free space closure (for example, ITGB1, KCNE3, and RASGRP1) did not show significantly altered cell proliferation in the c8161 melanoma cells (Fig. 2A). FN1 and FOS knockdowns in the c8161 melanoma cells showed significantly improved free space closure rates and both gene knockdowns also exhibited significantly reduced cell proliferation (Fig. 2A; rows 2-3 and columns 1-2). Likewise, no gene knockdowns in the c8161 melanoma cell line data set were associated with a reduced rate of free space closure and enhanced cell proliferation, or both an enhanced rate of free space closure and increased cell proliferation (Fig. 2A). In contrast, cell proliferation and cell motility in the HT1080 fibrosarcoma cells were impacted, with a reduction of cell proliferation the more pronounced effect. Together, these data suggest that in c8161 melanoma cells the set of gene knockdowns primarily affected free space closure through changes in cell motility, rather than through cell proliferation. The gene knockdowns influenced both cell motility and cell proliferation in the HT1080 fibrosarcoma cells but were more strongly associated with reduced cell proliferation.

### c8161 melanoma cells exhibit collective cell migration with a leading edge subpopulation that rapidly migrates in a directed manner to fill up open free space, regardless of gene knockdown

The preceding statistical analysis of open free space areas and cell count fold changes suggested that invasion is driven by cell motility in the c8161 melanoma cells but did not precisely identify which changes in motility at the cell-level contributed to reduced or enhanced free space closure. To address this, we performed cell tracking to investigate the motility properties of individual cells more thoroughly.

NC cells are well-known for their leader-follower dynamics, whereby leader cells must invade through immature ECM, loosely connected mesoderm and other cell types to pave the way for follower cells to move collectively behind the wavefront (Kulesa et al., 2021). To better understand whether the gene knockdowns in the screen perturb this pattern, we tracked c8161 melanoma and HT1080 fibrosarcoma cell trajectories throughout the free space closure process (Fig. 2B-H). In each of the wells, we tracked all cells in the field of view and computed the average displacement per frame for each cell. By making a distinction between cells at the leading (edge) versus cells in the follower (bulk) subpopulation, we computed the displacement difference between edge and bulk cells for each of the different wells, defined as the difference between the mean edge and the mean bulk cell displacement.

The displacement difference acts as a measure for how motile on average the cells at the edge of the collective are relative to those in the bulk subpopulation. A large displacement difference would indicate a highly active edge subpopulation that rapidly invades towards the center of the free space, with a less actively motile bulk population. Similarly, we computed the average directionality in the direction of the free space for each cell and determined the directionality difference as the difference between the average directionality for cells at the edge and that for cells in the bulk (Fig. 2B, inset). We find that c8161 melanoma cell edge subpopulations display significant differences in motility and directionality compared to the HT1080 fibrosarcoma cells across the different gene knockdowns (Fig. 2C, D). This finding suggests that c8161 melanoma but not non-NC-derived HT1080 cells exhibit an edge subpopulation that rapidly migrates in a directed manner to fill up free space, regardless of gene knockdown.

We next compared the displacement difference directly to free space closure (Fig. 2E-F). We find that for the c8161 melanoma cells, an edge subpopulation with enhanced motility leads to enhanced free space closure (Fig. 2G; Spearman correlation coefficient of −0.452, p-value=0.00180); a trend not observed in HT1080 fibrosarcoma cells (Fig. 2H). Together, this suggests that for c8161 melanoma cells the active leader cells within the edge subpopulation are most successful in closing the free space, in contrast to when all cells in the collective migrate according to the same rules. Lastly, the same analysis for the directionality difference showed no direct relationship with closure of the free space for the HT1080 fibrosarcoma cells.

### Leader cell motility is impaired after knockdown of BMP4, ITGB1, KCNE3, or RASGRP1 with only BMP4 knockdown also exhibiting less-directed, more wandering c8161 melanoma cell behaviors

Since our cell trajectory analysis suggested that highly motile leader cells are associated with faster free space closure (Fig. 2E; c8161 melanoma cells), we examined whether the set of gene knockdowns that result in reduced free space closure also exhibited a reduction in relative edge displacements. Indeed, leader c8161 melanoma cell motility is reduced by as much as 30% in the BMP4, ITGB1, KCNE3 and RASGRP1 knockdowns, with a mean cell motility of 5.1, 4.4, 4.5, and 5.2 um/hr, respectively, compared to 6.4 um/hr in controls (Fig. 2G). Furthermore, by characterizing the difference between displacement for leader versus bulk cells, we find that BMP4, ITGB1, and RASGRP1 knockdowns also exhibit a marked, reduced difference between leader and bulk cell motility. This was evident by the overlap in the inter-quartile range, contrary to the trend in control and KCNE3 (Fig. 2E). Interestingly, only BMP4 exhibited a reduction of leader cell directionality (0.094 versus 0.23 for control (Fig. 2H)), establishing that loss of BMP4 gives rise to less-directed, more wandering c8161 melanoma cell behaviors at the leading edge (Fig. 2H).

Performing this analysis on the remaining 10 out of the 14 gene knockdowns in c8161 melanoma cells that result in reduced free space closure (Fig. 2A; not significant in HT1080 fibrosarcoma cells), we find that all 10 genes except PRNP have significantly reduced edge cell displacement. No gene knockdown showed altered edge directionality as compared to the controls. These findings suggest that the cluster of gene knockdowns exhibited reduced free space closure due to inhibited leader cell motility. Furthermore, BMP4, ITGB1, and RASGRP1 also show a loss of leader cell-type behavior by displaying a larger difference between leader and follower cell motility. Finally, we remark that both FN1 and FOS knockdowns (which have significantly higher rates of free space closure than control) show the same trend, exhibiting significantly more motile edge cells (data not shown).

### Deep attention network analysis identified distinct cell-cell interaction patterns in collective cell migration of c8161 versus HT1080 cells and alterations after gene knockdown

We observed above that individual cell behaviors at the leading edge have a significant impact on the rate of free space closure, but cells do not migrate in isolation; closure is an inherently cooperative process. Therefore, we asked whether any of the 45 genes in the high-throughput screen play a role in cell-cell communication during the closure process. However, the analysis of large-scale cell trajectory data characteristic of high-throughput assays is incredibly time consuming.

Deep attention networks are a mathematical framework that enable quantitative measurement of the cell-cell interactions that most influence the velocity alignment in a collective group of cells. We trained the attention networks on each of the different siRNA gene knockdown data sets to quantify cell-cell attention patterns. These analyses show that cells ‘pay attention’ to a subset of neighbors such that only cells within a given region relative to the polarity axis of the cell influences the future direction of the cell (Fig. 3A). We first established typical characteristic patterns of ‘attention’ for c8161 melanoma and HT1080 fibrosarcoma cells in the absence of gene knockdown (Fig. 3B-C). In this view, c8161 melanoma cells are characterized by a highly localized attention pattern at the front of the cell. In stark contrast, we find a very narrow front-and-back pattern of attention for the HT1080 fibrosarcoma cells (Fig. 3C). This pattern is consistent with the typical sliding of cells in between each other in a largely non-coordinated fashion. Having characterized the baseline cell-cell attention patterns, we next turned to the question of whether the knockdown of any of the 45 genes would reveal attention patterns distinct from the wildtype scenario.

**Figure 3.**
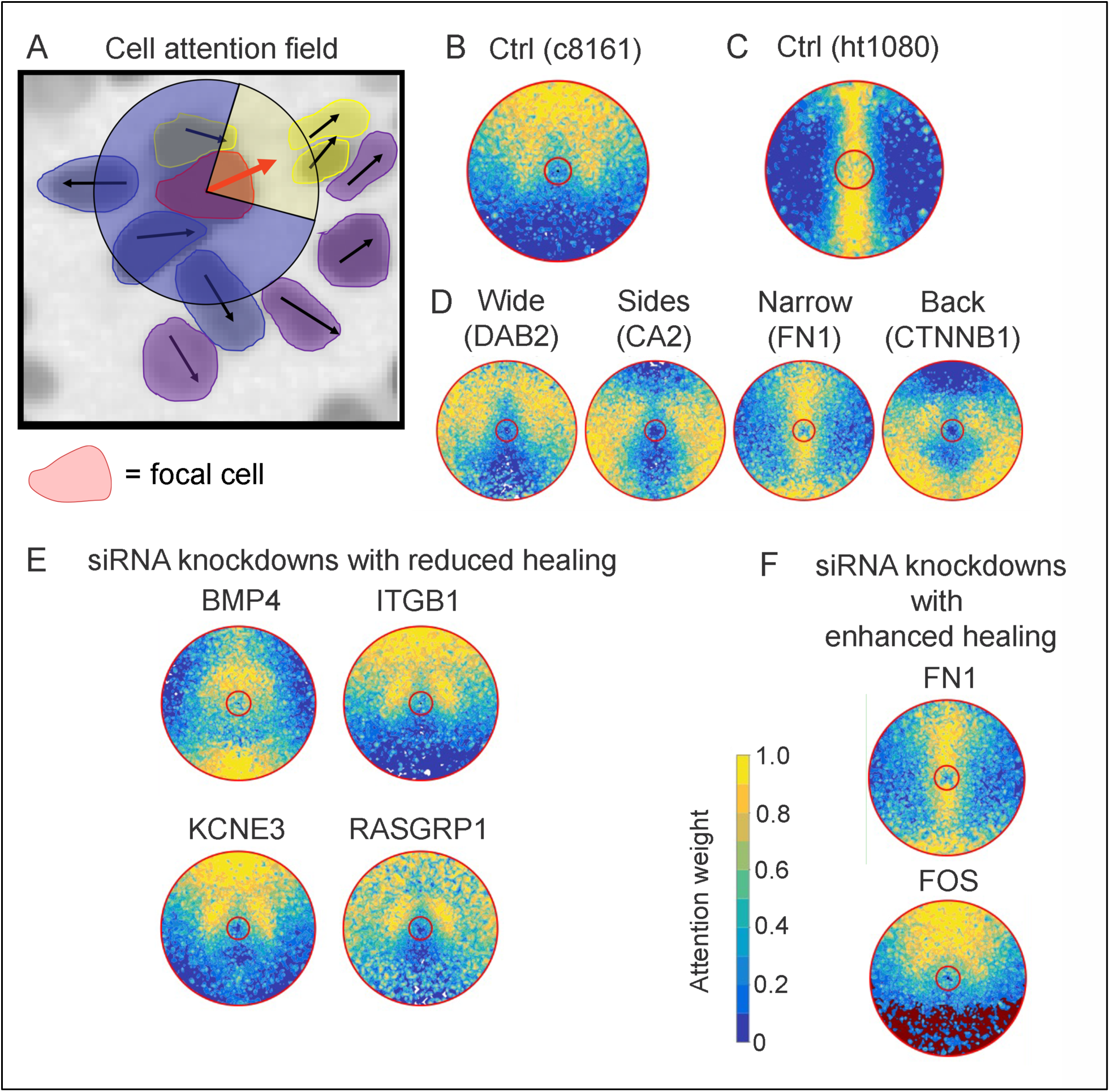
Deep learning network analysis reveals altered patterns of attention between cells. (A) Schematic showing focal cell (red) with a cartoon attention cone. Cells in yellow are cells in the high attention region, cells in blue in the low attention region and cells in green are outside the attention range. (B-C) Attention patterns for Neg knockdown (B) in c8161 and (C) in HT1080 cells. (D) Effects of siRNA knockdown on attention network patterns, notably the width of the attention cone as well as the location to the front or back of the cell has a wide variability. (E-F) Attention networks for (E) four siRNA knockdown perturbations that significantly affected wound closure rates and for (F) two siRNA knockdowns that showed enhanced healing.

To better compare patterns of cell neighbor relationships between the different gene knockdowns, we extracted quantitative metrics that describe the patterns of attention. We quantified both the extent to which cells pay attention to neighbors in front and behind as well as the width of the attention field. The c8161 melanoma cells have a much wider attention pattern than HT1080 fibrosarcoma cells (Fig. 3B, C). In addition, c8161 melanoma cells displayed a wide range of attention behaviors depending on gene knockdown (Fig. 3D-E). In particular, both the width of the cell-cell attention field as well as the front-to-back cell neighbor interactions were affected (Fig. 3D). For example, we observed a wider attention range in the DAB2 knockdown that stretched from the front to along the sides of the cell, suggesting the cell-cell interactions were broadened to side-by-side neighbors (Fig. 3D). In contrast, the knockdown of CA2 produced a pattern where regions ahead and behind the cell were largely ignored (Fig. 3D). FN1 knockdown produced a pattern in the c8161 melanoma cells that resembled the wildtype HT1080 cells with a front-to-back pattern (compare Fig. 3C with Fig. 3D). In the CTNNB1 knockdown, cells appeared to pay attention to neighbors along the sides and behind rather than search for open free space (compare Fig. 3B and Fig. 3D). Strikingly, in the HT1080 fibrosarcoma cells we find that there is no qualitative variation in attention patterns (data not shown), consistent with the observation that migration is highly collective in c8161 melanoma cell populations, but not in the HT1080 fibrosarcoma cell populations.

We next asked whether altered patterns of attention are associated with the observed free space closure changes. We compared the attention patterns for the four gene knockdowns leading to inhibited free space closure (BMP4, ITGB1, KCN3, RASGRP1) with the attention patterns for the knockdowns that resulted in improved free space closure (Fig. 3E). While loss of ITGB1 and KCNE3 retained the wildtype pattern of attention, both BMP4 and RASGRP1 knockdowns displayed altered attention patterns (Fig. 3E). Importantly, these respective attention patterns were qualitatively different; loss of BMP4 displayed attention towards the front and back and loss of RASGRP1 showed attention only towards the front (Fig. 3E). Likewise, loss of FN1 and FOS promoted free space closure but showed no common qualitative features in the attention pattern (Fig. 3E).

### Addition of recombinant proteins for five secreted genes of the 45-gene panel increases migration of c8161 melanoma cells in culture

Eleven of the genes from the 45-gene panel are known secreted molecules. To test whether the presence of any of these factors stimulate cell migration, we added recombinant proteins for BMP4, DKK3, ESM1, FN1, IFI30, Ly86, PNRP, SERPINI1, UPP1, VEGFC separately to the media with cultured c8161 melanoma or HT1080 cells in the plugged plate assays (Fig. 4A-B). After addition of recombinant proteins, rubber stopper plugs were removed and measurements of cell positions revealed that BMP4, ESM1, FN1, IFI30, and VEGFC stimulate cell migration, resulting in a significant increase of the area of open free space covered by the invading c8161 melanoma cells (Fig. 4C). Addition of SERPINI1 led to a similar observation in c8161 cells but the change was only significant in HT1080 fibrosarcoma cells (Fig. 4C). High c8161 melanoma cell density in culture was also observed after addition of exogenous protein for UPP1, without increased cell migration (data not shown). A decrease in uniform density with increased clustering was observed in c8161 melanoma cells after addition of DKK3 (Fig. 4B-C).

**Figure 4.**
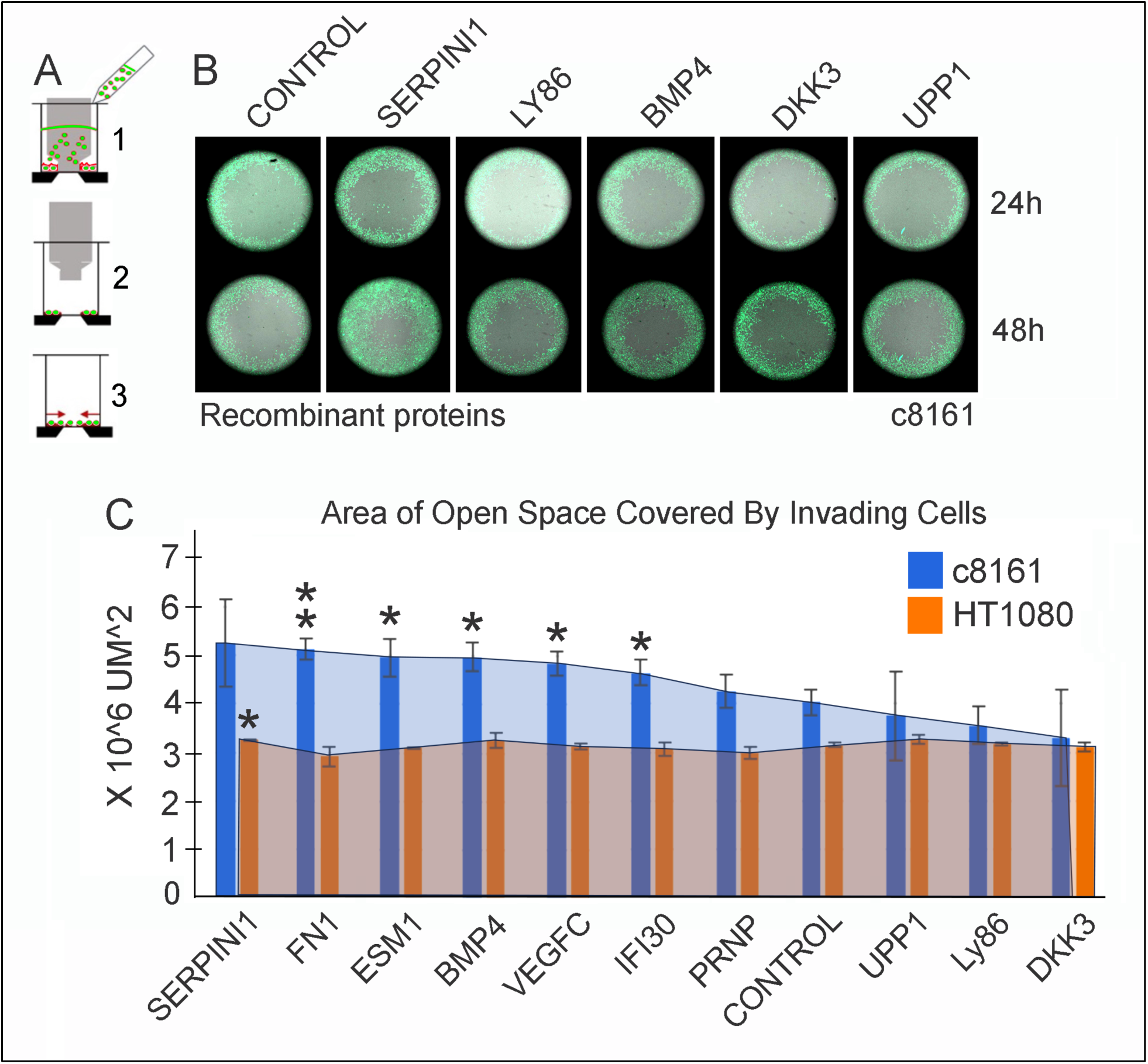
Recombinant proteins affect migration of human c8161 melanoma cells into open free space in culture. (A) c8161 cells were seeded into plugged 96-well plates and plugs were removed after 24 hours; recombinant proteins were added to the culture media. Cell plates were imaged at +24 hours and at 48 hours after plug removal, and cell positions were recorded and scored. (B) Typical examples of c8161 melanoma cell positions at 24 hours (top row) and 48 hours (bottom row) after plug removal and addition of recombinant protein. (C) Compared to control (PBS), c8161 cells in the presence of VEGFC (p=0.0186), BMP4 (p=0.0134), ESM1 (p=0.0262), IFI30 (p=0.0469), and FN1 (p=0.0049) significantly increased migration into the open free spaces. SERPINI1 was the only recombinant protein to affect migration of HT1080 cells (p=0.0451). All recombinant proteins were used at 50ng/ml added to the culture media. c8161 cells (blue bars) and HT1080 cells (orange bars) with p-values calculated as * = p<0.05, ** = p<0.01.

### Knockdown of BMP4 by siRNA inhibits *in vivo* c8161 melanoma cell invasion in a chick embryo transplant model, but invasion may be rescued *in vitro* after addition of BMP4 recombinant protein

The discovery that several recombinant proteins of the 45-gene panel stimulate c8161 melanoma cell invasion/migration suggested synergy experiments to rescue invasive cell behaviors after gene knockdown. We knocked down the function of BMP4 by siRNA in c8161 cells and loaded cells into the rubber stopper assay (Fig. 5A). After 24 hours, BMP recombinant protein or control (PBS) was added to the culture media and the rubber stopper barrier was removed (Fig. 5A). Measurements of the area covered by c8161 melanoma cells showed a significant decrease after only BMP4 siRNA knockdown (PBS control, Fig. 5B). However, the addition of BMP4 to the culture media stimulated c8161 melanoma cell migration so that nearly double the area was covered by invasive cells even with BMP4 siRNA targeted to the cells (Fig. 5B). To test whether knockdown of BMP4 inhibits *in vivo* cell invasion/migration, we transplanted c8161 melanoma cells treated with BMP4 siRNA knockdown and grown in a hanging drop into the chick hindbrain (at the level of approximately r4) or adjacent to r4 (Fig. 5C). We find that c8161 melanoma cells transplanted into the chick neural tube became significantly less invasive (Fig. 5C), with slightly lesser effects in cells transplanted adjacent to the hindbrain (Fig. 5C). Treatment of the c8161 melanoma cells with Noggin (an antagonist to BMP4) showed a significant reduction in cell invasion/migration from the chick neural tube (Fig. 5C).

**Figure 5.**
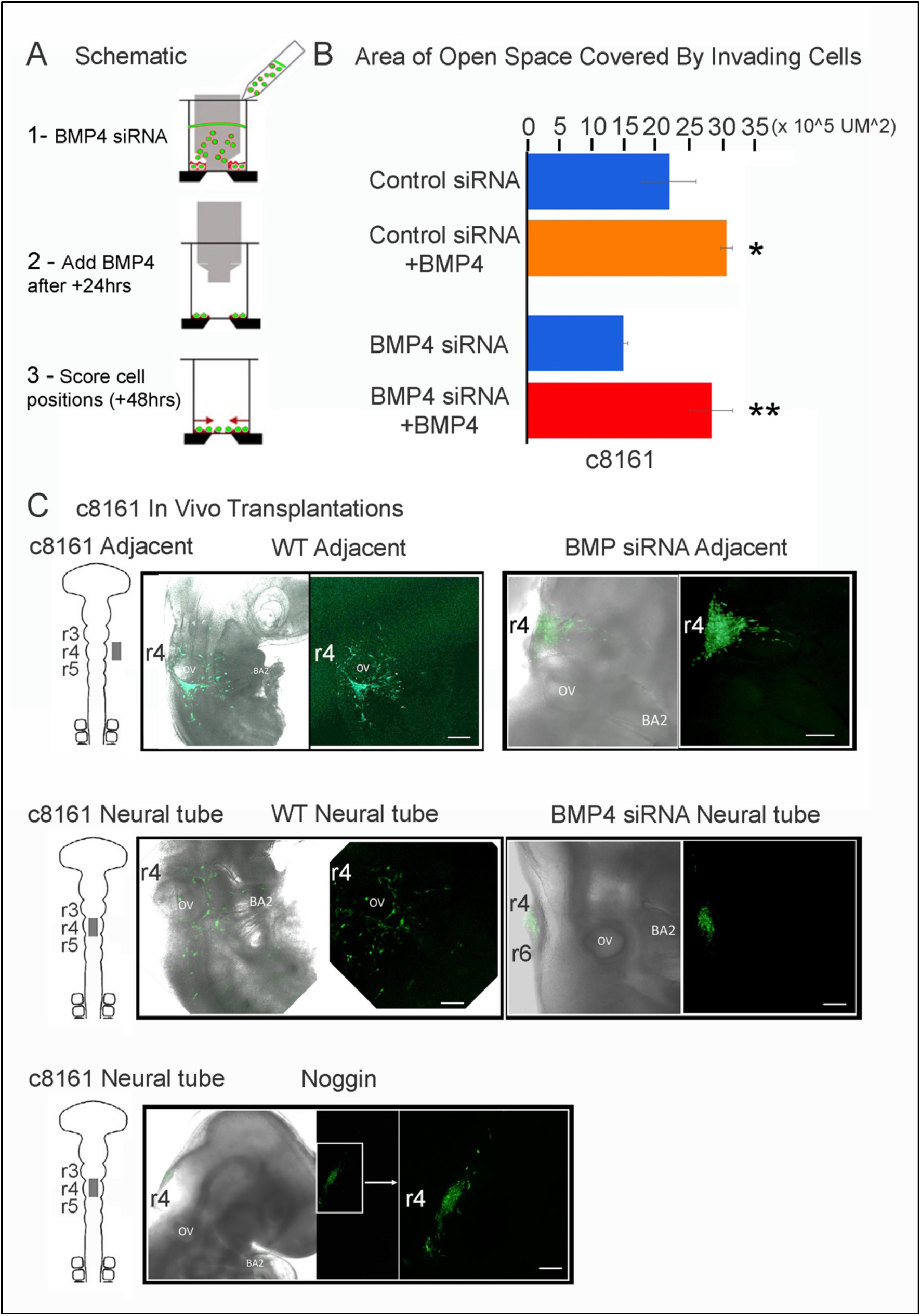
Exogenous BMP4 rescues BMP4 siRNA knockdown in human c8161 melanoma cells in culture. (A) c8161 cells were seeded into plugged 96-well plates after BMP4 siRNA knockdown and plugs were removed after 24 hours; BMP4 recombinant protein was then added to the culture media. Cell plates were imaged at +24 hours after plug removal and cell positions were recorded and scored. (B) The area covered by invading c8161 melanoma cells was measured and compared for four scenarios including control siRNA, control siRNA+BMP4, BMP4 siRNA, and BMP4 siRNA+ BMP4. (C) Typical examples of c8161 melanoma cell positions after knockdown of BMP4 by siRNA or Noggin treatment and transplantation into the chick hindbrain (∼r4 level) or the adjacent paraxial mesoderm and three-dimensional confocal images collected after + 24 hours. The BMP4 recombinant protein was added to the culture media at 50ng/ml. p-values calculated as * = p<0.05, ** = p<0.01.

## DISCUSSION

Previous single-cell profiling of chick cranial NC cell migratory streams provided a basis to compare a novel transcriptional signature of the migratory stream wavefront with gene signatures obtained from a broad range of other cell invasion phenomena (Morrison et al., 2021). This produced a 45-gene panel too time-consuming and impractical for single gene *in vivo* functional studies in the chick embryo, but appropriate for an siRNA high-throughput screening assay. Comparison of changes in cell migration into the open free space and cell number after each of 45 siRNA gene knockdowns in NC-derived human c8161 metastatic melanoma cells versus non-NC-derived HT1080 fibrosarcoma cells, converged on a subset of 14 out of 45 genes that significantly impact cell invasion. Confocal time-lapse imaging provided a very large set of cell trajectory data sufficient for statistical analysis and deep learning attention network analysis of cell neighbor relationships of both leading (edge) and follower (bulk) subpopulations. Gain-of-function experiments with 11 known secreted molecules of the 45-gene panel identified six genes that enhanced c8161 melanoma cell migration when the relevant recombinant proteins were added to the culture media.

These top six out of the 14 genes we identified from the 45-gene panel whose knockdown significantly affected free space closure did so with changes in c8161 melanoma cell motility rather than cell proliferation. Key to accurately making this distinction was the use of a rubber stopper in the high- throughput assay which overcame the problems of rupturing cell membranes at the wound edge and creating a non-uniform invasive wavefront, typical of a scratch assay. This allowed for a more accurate means to collect precise cell trajectory data of an invasive migratory wavefront and distinguish leading edge versus bulk follower cell movements. The top six out of 14 genes whose knockdown resulted in significantly slower free space closure were BMP4, ITGB1, IFI30, KCNE3, PRKCQ, and RASGRP1. RASGRP1 (a Ras guanine nucleotide-releasing factor that activates Ras and its downstream ERK and AKT signaling pathways) has been shown to promote the formation of spontaneous skin tumors and enhance malignant progression in a multi-stage carcinogenesis skin model that relies on oncogenic activation of H-Ras (Sharma et al., 2014). Similarly, there is previous evidence to support a predominant role for BMP4 in melanoma metastasis, in part through regulation of SNAI2 (Gupta et al., 2005; Rothhammer et al., 2005). More recent studies have shown that BMP4 enhances the epithelial-to-mesenchymal transition (EMT) and cancer stem cell properties of breast cancer cells via Notch signaling (Choi et al., 2019). However, the role of BMP4 in cell invasion is less clear (Ehata and Miyazono, 2022). Interestingly, activation of BMP signaling in SMAD-4 negative colorectal cancer cell lines increased cell migration, invasion and formation of invadopodia (Voorneveld et al., 2014), while BMP inhibitors show promise at reducing melanoma growth in mouse melanoma models (Kalal et al., 2021). Together, this encourages further examination of BMP4 in NC cell migration. In support of this, we previously observed high induction of BMP4 in c8161 melanoma cells at the migratory wavefront after cell transplantation into the chick embryonic microenvironment (Bailey et al., 2012). Thus, further analysis of NC cell behaviors *in vitro* and *in vivo* in the chick embryo transplant model after loss of BMP4, RASGFP1 and the other four genes may provide insights into their functional roles in cranial NC cell migration.

Deep learning attention network analysis identified two key aspects of cell-cell neighbor interactions from the high-throughput assay cell trajectory data. First, there were clear differences in collective attention between the c8161 melanoma versus HT1080 fibrosarcoma cells -- the latter was used as a negative non-neural crest derived cell line control (Fig. 3). For the HT1080 cells, the network analysis determined that the most influential neighbors were the leader directly in front of and follower directly behind the focal cell (Fig. 3). This was somewhat expected given the known cell behavior of HT1080 cells moving in strongly aligned chains of cells in a leader-follower manner.

In contrast, the collective attention network for c8161 melanoma cells determined the most influential neighbors to be within a wide front ahead of and to the sides of the focal cell (Fig. 3). This analysis is consistent with observations of the collective migratory behaviors of the c8161 melanoma cells, edge versus bulk subpopulations (Fig. 3). Second, in several cases in which one of the 45 gene panel was knocked down, there was a wide range of significant changes to the c8161 melanoma cell wildtype attention map, including an expansion of the cell-cell neighbor interactions along the sides and to the rear of the focal cell, and even a switch to the follow the leader directly in front pattern (Fig. 3). When we asked whether altered patterns of attention were associated with the observed free space closure changes for the top four gene knockdowns leading to significantly reduced free space closure (BMP4, ITGB1, KCN3, RASGRP1), we learned that only BMP4 and RASGRP1 knockdowns displayed altered attention patterns (Fig. 3E). Loss of BMP4 displayed a reduced forward attention map and a long-range connection with cells in the rearward direction, including a curious gap directly behind the focal cell (Fig. 3). In contrast, loss of RASGRP1 showed an expanded attention map to the front and sides of the focal cell (Fig. 3), and loss of ITGB1 and KCNE3 retained the wildtype c8161 melanoma cell attention map (Fig. 3D). Future studies that fluorescently label cell membranes may be able to distinguish and measure changes in cell-cell contacts between neighbors that would enable further interpretation of the deep learning attention map results.

We also unexpectedly learned of major alterations to the cell attention maps after loss-of-function of DAB2, CA2, FN1, and CTNNB1; genes that did not correlate with significant reduction in free space closure with the exception of FN1 (Fig. 3D). For example, after loss of DAB2, cells appeared to expand communication with neighbors along the sides of the cell rather than simply in front of the cell in search of open free space. After loss of FN1, the alteration of cell-cell interactions was more dramatic. Loss of FN1 led to a front-to-back cell-cell interaction pattern, suggestive of a less coordinated sliding of cells between each other, similar to the control pattern of HT1080 cells (Fig. 3). DAB2, which encodes a mitogen-responsive phosphoprotein, has been shown to regulate the migration and invasion of prostate cancer cells (Westcott et al., 2015). Its knockdown by shRNA inhibits these characteristics in PC3 cells *in vitro*, but the mechanism of its action is unclear. In our hands, when DAB2 was knocked down by siRNA, c8161 melanoma cells altered their cell neighbor relationships to include a much wider attention field (Fig. 3) but did not show a significant reduction in free space closure *in vitro*. These observations suggest that either DAB2 is not critical for melanoma migration or its knockdown causes cells to seek alternative guidance signals.

The addition of recombinant proteins into the culture media to rescue migration of siRNA-treated c8161 melanoma cells identified a role for BMP4 and directed future gene synergy experiments. The addition of recombinant proteins into the culture media of c8161 melanoma and HT1080 fibrosarcoma cells for 10 out of the 11 known secreted molecules of the 45-gene panel (Fig. 4A, B) identified five out of 10 molecules (BMP4, ESM1, FN1, IFI30, VEGFC) that stimulate c8161 cell migration and significant increases in invasion of the open free space area (Fig. 4C). Either BMP4 knockdown by siRNA or Noggin significantly reduced *in vivo* invasion/migration after transplantation into the chick embryo hindbrain (Fig. 5B, C). However, addition of BMP recombinant protein into the culture media, 24 hours after siRNA knockdown, stimulated migration and increased the covered free space area (Fig. 4C). SERPINI1 showed a significant change in open free space area with HT10180 cells but was not significant in c8161 cells (Fig. 4C). These data support observations made in the colorectal cancer cell line data mentioned above (Voorneveld et al., 2014; Kalal et al., 2021) and motivate potential future synergy experiments in which any of the other four recombinant proteins may be added after siRNA knockdown and rubber stopper removal of the other 14 out of 45 candidates that reduced c8161 melanoma cell migration after siRNA knockdown.

In summary, our study demonstrated the rapid and systematic analysis of a 45-gene panel to converge to on a critical subset of genes using a combined experimental, statistical, and deep learning approach. As advances in spatial transcriptomics and single cell profiling bring us closer to identifying gene sets associated with distinct, dynamic biological processes, so too will upgrades in deep learning and AI algorithms that shorten the time between high-throughput screening and single gene functional analysis. Deep learning network analysis provided novel insights, but it is an open question to how to interpret these attention maps to provide mechanistic insights. In the chick cranial NC model system, it will be interesting to now determine whether the function of the identified critical subset of cell invasion genes is conserved for other NC cell migratory streams throughout the head, heart and trunk axial levels and in other vertebrate model organisms. Further understanding of the function of NC cell invasion genes will have direct applications to emerging cell replacement therapies to repair neurocristopathies by providing a molecular blueprint to control transplanted cell behaviors and suggest therapeutic applications to inhibit NC cell-derived cancer metastasis.

## EXPERIMENTAL PROCEDURES

c8161 human malignant aggressive melanoma cells were provided by Dr. Mary Hendrix (Children’s Memorial Research Center, Chicago, IL) and were maintained in RPMI media supplemented with 10% FBS. HT1080 cells were obtained from ATCC (catalog #CCL-121) and maintained in EMEM media supplemented with 10% FBS. Cells were plated in Oris collagen coated 96-well plugged assay plates (Platypus Technologies, #CMACC5.101). HT1080 cells were seeded at 35,000 cells/well and c8161 cells at 45,000 cells/well. For siRNA knockdown, three siRNAs targeting each were purchased from Ambion (Silencer Select, standard purification). Cell lines were transfected with siRNAs using xTremeGene siRNA transfection reagent (1ul/well, Roche, #4476093001; standard protocol). After addition of siRNA transfection mix, 96-well plugged plates were incubated for 48 hours (37C, 5% CO2). Plugs were then removed with the provided plug removal tool (Oris) and 100ul of fresh media added to each well. Plates (with lid on) were placed in the Opera Phenix High-Content Screening System (Perkin Elmer; prewarmed to 37C and 5% CO2). Images were collected every 30 min for 24 hours.

### Quantitative analysis

#### Cell count and wound areas

For the computation of wound areas, we used Fiji to threshold and binarize the nuclear channel to obtain the individual cell locations, and then performed convolution with a Gaussian kernel with standard deviation of two cell widths to smooth the nuclear channel into a density field. We then thresholded and binarized to obtain wound masks to measure the area of the wound. We used the inbuilt Laplacian of Gaussian (LoG) detector in Fiji to detect spots from the nuclear channel to count cells using Fiji and obtain cell count fold changes between the start and the end of the experiment.

#### Statistical analysis of fold changes

We computed the area and cell count fold changes for each of the tissues recorded and group them together by knockdown, giving sets of 3-9 biological samples for each individual knock-down. We computed z-scores for each of the wound area and cell count fold changes and performed a standard Welch’s t-test to compute the corresponding p-values for wound area and cell count fold change.

#### Individual cell trajectories

We use TrackMate in Fiji to obtain individual cell tracks from the binarized and thresholded images. We use the inbuilt LoG detector with standard LAP tracker in Fiji/Trackmate with no track splitting and merging, as well as a gap closure of five frames and a maximum linking distance of 100μm. We set the cell diameter for c8161 cells to 20μm and that for ht1080 cells to 30 ums. From the TrackMate outputs, we filtered for cell tracks that contain at least five consecutive frames.

#### Classification of edge and bulk cells

We classified cells as edge cells if their distance to the masked wound edge was less than five cell radii at their first detected time point.

#### Training of deep attention networks

Training was done with the same hyperparameters as Lachance et al.(2022). Training was performed on the Oxford Advanced Research Computing cluster using five 48-core Cascade Lake (Intel Xeon Platinum 8268 CPU @ 2.90GHz) nodes.

#### Data availability

Trained networks and their inputs are available at the Zenodo repository.

## ACKNOWLEDGEMENTS

PMK would like to thank the generous funding of NIH/NICHD (R03-HD105079-02) for partial support of this study. JCK would like the thank the generous support of the Stowers Institute and core facilities, including Dr. Jay Unruh.

